# A persistent lack of International representation on editorial boards in environmental biology

**DOI:** 10.1101/131292

**Authors:** Johanna Espin, Sebastian Palmas, Farah Carrasco-Rueda, Kristina Riemer, Pablo E. Allen, Nathan Berkebile, Kirsten A. Hecht, Kay Kastner-Wilcox, Mauricio M. Núñez-Regueiro, Candice Prince, Constanza Rios, Erica Ross, Bhagatveer Sangha, Tia Tyler, Judit Ungvari-Martin, Mariana Villegas, Tara T. Cataldo, Emilio M. Bruna

**Author notes:** Current Address: Smithsonian Migratory Bird Center, Washington, DC, USA. Current Address: School of Natural Resources & Environment, University of Florida, Gainesville, USA.

## Abstract

The scholars comprising journal editorial boards play a critical role in defining the trajectory of knowledge in their field. Nevertheless, studies of editorial board composition remain rare, especially those focusing on journals publishing research in the increasingly globalized fields of science, technology, engineering, and math (STEM). Using metrics for quantifying the diversity of ecological communities, we quantified international representation on the 1985-2014 editorial boards of twenty-four environmental biology journals. Over the course of three decades there were 3831 unique scientists based in 70 countries that served as editors. The size of the editorial community increased over time – there were 420% more editors serving in 2014 than in 1985 – as did the number of countries in which editors were based. Nevertheless, editors based outside the ‘Global North’ (the group of economically developed countries with high per capita Gross Domestic Product (GDP) that collectively concentrate most global wealth) were extremely rare. Furthermore, 67.06% of all editors were based in either the USA or UK. Consequently, Geographic Diversity – already low in 1985 – remained unchanged through 2014. We argue that this limited geographic diversity can detrimentally affect the creativity of scholarship published in journals, the progress and direction of research, the composition of the STEM workforce, and the development of science in Latin America, Africa, the Middle East, and much of Asia (i.e., the ‘Global South’).

## INTRODUCTION

There are currently over 28,000 peer-reviewed academic journals [1], and the scholars who serve on the editorial boards of these journals play a major role in defining the trajectory and boundaries of knowledge in their disciplines [2]. This is because board members are responsible for coordinating the evaluation by outside experts of a manuscript’s technical aspects and the “importance” or “novelty” of the research it summarizes, i.e., peer review, on which the decision to publish a manuscript is ultimately based. Editors also play a central but underappreciated role in shaping the community of scholars contributing to the discourse in their field. First, by recommending the publication of an article the editor confers legitimacy not only on the research, but also upon the individuals who carried it out [3,4]. Second, editors help choose new editors. In doing so, they confer enhanced status and visibility on a select group of scholars who then benefit from the unique opportunities for professional advancement provided by board membership [5]. Editors are therefore a small but powerful group of “Gatekeepers” [2] that select the scientists and ideas shaping the direction of their discipline.

The increased recognition of editor power, along with the results of studies on workforce diversity [6], have heightened concerns about how the composition of editorial boards might influence the peer-review process [7]. For example, it has been suggested that boards whose members are demographically homogenous might converge on a narrow suite of research topics and approaches they consider worthy of publication [3,4]. This narrow vision – and the board structure driving it – could be perpetuated by editors nominating collaborators, whose perspectives and backgrounds likely match their own, for board service. Indeed, this is among the principal reasons put forward to explain why women remain severely underrepresented on editorial boards across academic fields [5], which has consequences for the selection of referees and other critical aspects of the editorial process [8].

Recent decades have seen the rapid globalization of research in science, technology, engineering, and math (STEM), resulting in greater representation in international journals of authors based in the ‘Global South’ [9,10]. i.e., the world’s ‘developing’ or ‘emerging’ economies located primarily in Latin America, Asia, Africa, and the Middle East. Having editorial boards that reflect this increasing ‘geographic diversity’ of the global scientific community is thought to benefit both journals and disciplines in ways that parallel the benefits resulting from other forms of diversity. In field-based sciences such as ecology or geology, for example, editors based in the region where studies are conducted will be more familiar with the environmental, social, and economic context and constraints under which they were carried out [11]. This could ensure both more rigorous review and a fairer assessment of reviewer criticisms and proposed improvements. Furthermore, scientists trained in different parts of the world can also have very different epistemological orientations. Increasing geographic diversity on an editorial board could therefore broaden the scope of theoretical and methodological approaches a journal publishes. Ultimately, these benefits of internationalization could help to minimize apparent biases in the review, publication, and citation of articles based on an author’s nationality or home-country [10,12].

The first systematic efforts to quantify the nationality of STEM editors – often by using the country in which they were based as a proxy for nationality – were carried out in the early 1980’s [13,14]. Since then a small but growing number of studies have observed similar patterns to what these early ones did – individual editorial boards tend to be dominated by scholars from or based in the United States of America (USA) and the United Kingdom (UK) [7]. However, prior studies typically compared board composition of journals using data from only a single calendar year, which makes it impossible to evaluate how the community of gatekeepers has changed over time. Furthermore, most of the journals reviewed are from the physical sciences, medical fields, or lab-based biological sciences [4,15]. As a result, almost nothing is known about geographic diversity of editors in field-based STEM disciplines [16] such as ecology, evolution, and natural resource management (hereafter “environmental biology”, EB).

The term “diversity” is often used colloquially to refer to the representation of different groups in a focal population or workplace. However, one can formally quantify the diversity of a community (e.g., an assemblage of editors) using a suite of indices derived from information theory. While the indices differ in their assumptions and applications, the most commonly used are calculated using two types of information: the number of categories found in a sample (i.e., “richness”) and the relative abundance of these categories (i.e., “evenness”). Most studies of editorial board composition to date only report the number of countries represented by editors, i.e., Geographic Richness. However, diversity indices permit a more nuanced evaluation of community composition. For example, using only Richness might lead one to conclude that the geographic representation of editors based in different countries has remained steady over time, when in fact one country has become numerically dominant. Another advantage of diversity indices is that they can be compared across groups (e.g., journals), even if the groups differ in richness or population size.

We identified all scientists serving from 1985-2014 on the editorial boards of 24 leading journals in Environmental Biology (Table S1) and the countries in which they were based during their board tenure. We then calculated the Geographic Richness and Geographic Diversity of this editor community and quantified how it has changed over the last three decades. Finally, we assessed the geographic distribution of editors at broader geographic and macroeconomic scales by comparing the representation of editors from different World Bank geographic regions and national income categories (for details on data collection and analysis see Text S1).

### How geographically diverse is the editorial community?

Between 1985-2014, N = 3831 scientists served as editors for our N = 24 focal journals. The size of the editor community increased steadily over time, with 420% more editors serving in 2014 than in 1985 (Fig 1A). Not surprisingly, this led to an increase in the Geographic Richness of the editor community – the number of countries represented by editors in 2014 was 52% higher than in 1985 (N=52 vs. N = 34), and the number of countries to have been represented by at least one editor increased from 34 to 70 (an increase of 86%; Fig 1B). However, scientists based in the USA and UK made up an overwhelming majority of the editor community (55.29% and 11.77%, respectively; Fig 2A). Although there have been modest increases (≤2%) from 1985 to 2014 in the proportion of editors based in five other countries (S1 Text Fig C), the continued concentration of editors in a very small number of countries is why the low-Geographic Diversity observed in 1985 has remained unchanged through 2014 (Fig 1C, Table A in S1 Text).

**Fig 1.**
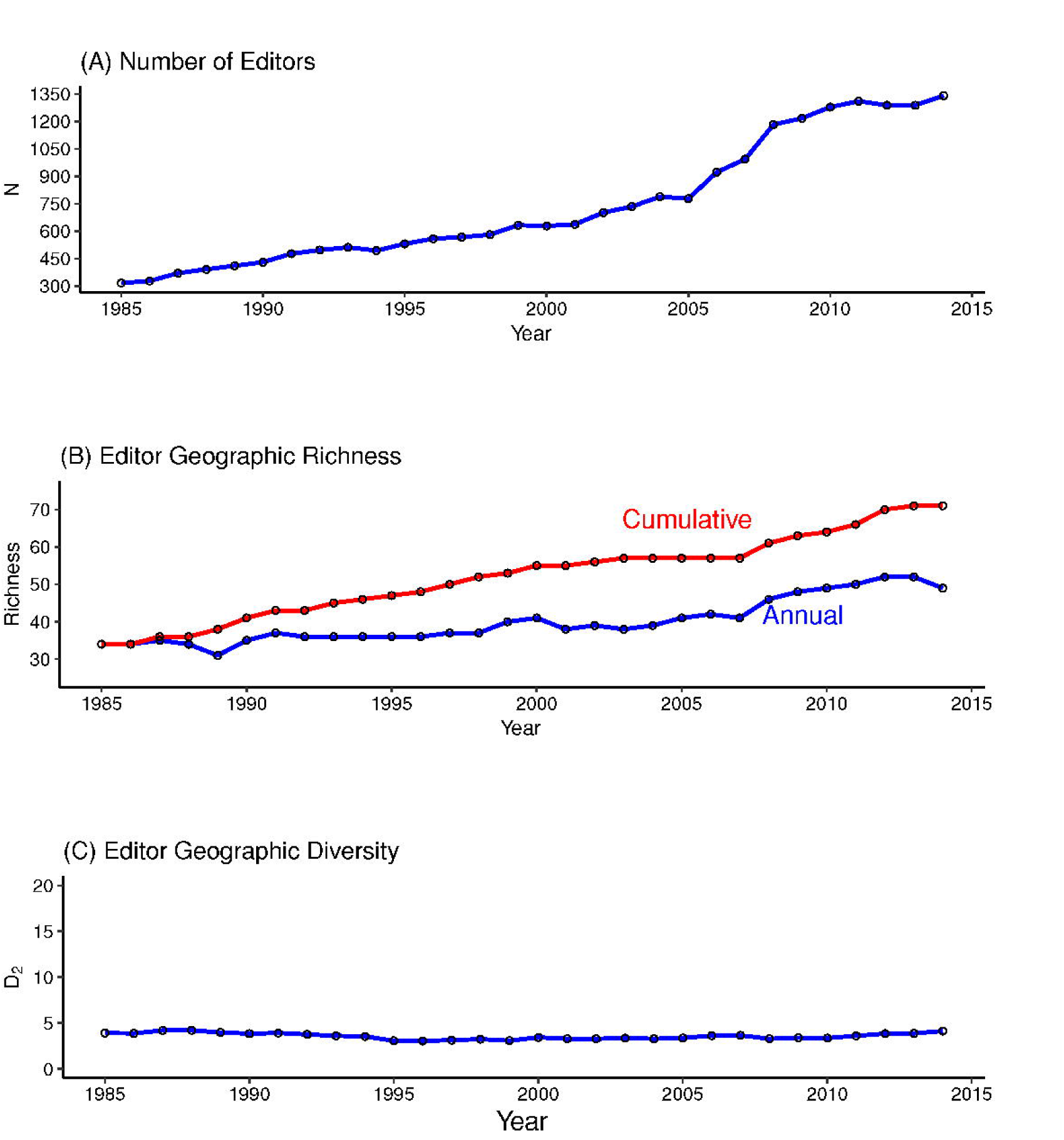
Community composition of editors in environmental biology (1985-2014). (A) Geographic Richness: Cumulative Richness is the total number of countries represented by at least one editor through a given year; Annual Richness is the total number of countries represented by editors in each year (B) The total number of unique editors serving each year from 1985-2014 (C) the Geographic Diversity of editors in environmental biology each year from 1985-2014. We measured diversity using the reciprocal of Simpson’s index, *D*_*2*_. Larger values of *D*_*2*_ indicate greater diversity, with the maximum potential diversity (MPD) equal to the greatest number of countries represented in any one year of the survey (MPD Editors = 52). For additional details see S1 Text.

**Fig 2.**
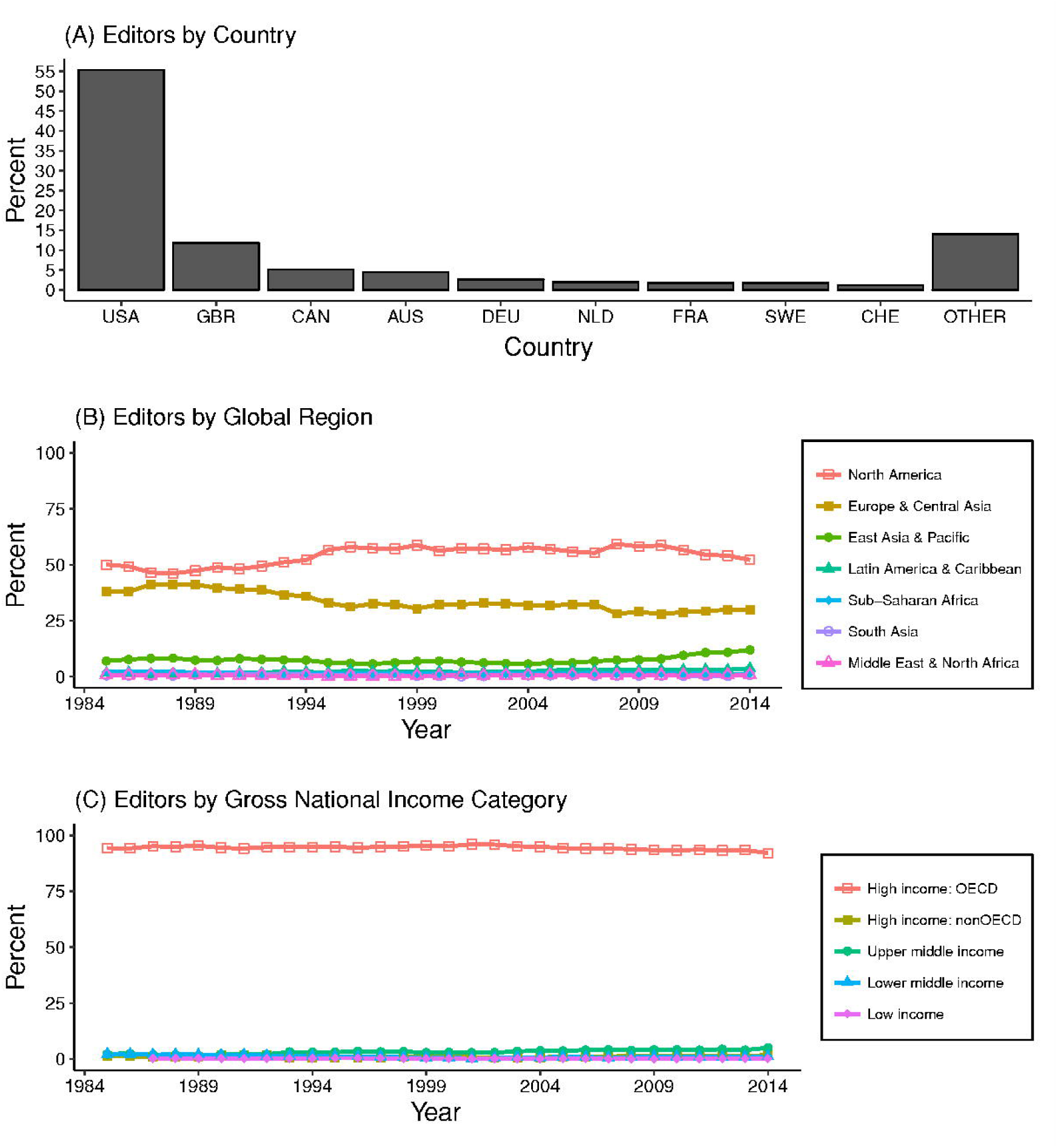
The percentage of environmental biology editors based in different countries, global regions, and World Bank national income categories. (A) Countries; Abbreviations: USA: United States of America, GBR: United Kingdom, CAN: Canada, AUS: Australia, NLD: Netherlands, FRA: France, SWE: Sweden, CHE: Switzerland. (B) World Bank global regions (C) World Bank Gross National Income categories.These patterns are echoed when assessing representation at broader geographic or macroeconomic scales. The proportion of editors each year that were based in North America varied from 46%-59%, while 28-41% were based in Europe/Central Asia (Fig 2B-C). The number of editors from the East Asia/Pacific region doubled from 1985 to 2014 (5.6% and 11.9%, respectively; Text S1 Fig B), but most of these were in the high-income countries of Australia, New Zealand, Singapore, and Japan. This concentration of editors in the Global North – the group of economically developed countries with high per capita Gross Domestic Product (GDP) that collectively concentrate most global wealth [17] – was observed at all levels of the gatekeeper hierarchy: 94% of Subject and Associate Editors, and a remarkable 98.2% of Editors-in-Chief, are based in high-income countries or Western Europe (Table 1, Text S1). In contrast, we found only a fraction of editors have been based in the Global South (Fig 2B-C). For example, Brazil, Mexico, and China are represented by fewer editors than Sweden, New Zealand, and the Netherlands (Number of Editors in 2014: Netherlands = 40, Sweden = 25, New Zealand = 26, China = 22, Brazil = 15, Mexico = 9).

**Table 1:**
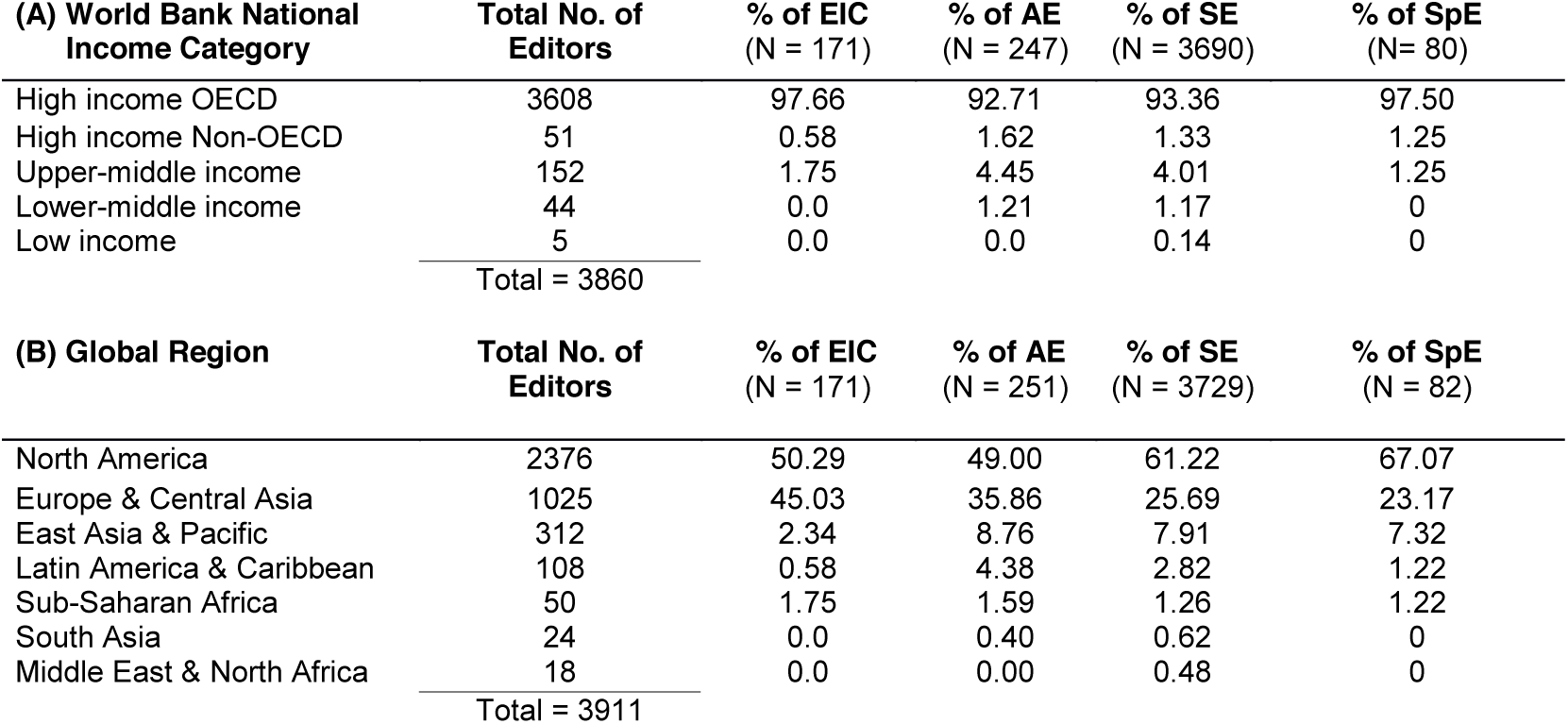
Percentage of the editorial board members from N = 24 environmental biology journals based in different (A) World Bank Country Income Categories and (B) Global Regions. Between 1985-2014 there were N = 3831 unique editors from 70 countries. The total number of editors in each region and national income category differs due to some editors having moved between 1984-2015; similarly, one person may serve multiple editorial roles. Numbers in parentheses are the number of unique editors in each category. Abbreviations: EIC: Editor-in-Chief, AE: Associate Editor, SE: Subject Editor, SpE: Special Category Editor.

| **(A) World Bank National Income Category** | **Total No. of Editors** | **% of EIC**<br>**(N = 171)** | **% of AE**<br>**(N = 247)** | **% of SE**<br>**(N = 3690)** | **% of SpE**<br>**(N = 80)** |
| --- | --- | --- | --- | --- | --- |
| High income OECD | 3608 | 97.66 | 92.71 | 93.36 | 97.50 |
| High income Non-OECD | 51 | 0.58 | 1.62 | 1.33 | 1.25 |
| Upper-middle income | 152 | 1.75 | 4.45 | 4.01 | 1.25 |
| Lower-middle income | 44 | 0.0 | 1.21 | 1.17 | 0 |
| Low income | 5 | 0.0 | 0.0 | 0.14 | 0 |
| Total = 3860 |  |  |  |  |  |
| **(B) Global Region** | **Total No. of Editors** | **% of EIC**<br>**(N = 171)** | **% of AE**<br>**(N = 251)** | **% of SE**<br>**(N = 3729)** | **% of SpE**<br>**(N = 82)** |
| North America | 2376 | 50.29 | 49.00 | 61.22 | 67.07 |
| Europe & Central Asia | 1025 | 45.03 | 35.86 | 25.69 | 23.17 |
| East Asia & Pacific | 312 | 2.34 | 8.76 | 7.91 | 7.32 |
| Latin America & Caribbean | 108 | 0.58 | 4.38 | 2.82 | 1.22 |
| Sub-Saharan Africa | 50 | 1.75 | 1.59 | 1.26 | 1.22 |
| South Asia | 24 | 0.0 | 0.40 | 0.62 | 0 |
| Middle East & North Africa | 18 | 0.0 | 0.00 | 0.48 | 0 |
| Total = 3911 |  |  |  |  |  |

Although several explanations have been put forward to account for this disparity, we believe one of the most common ones – a dearth of capable scientists in the Global South from which to draw [18] – is unlikely to be the cause. The number of scientists in the Global South is increasing dramatically, both in absolute terms and per-capita [19], as is their productivity [10,16,20]. Therefore, the number of scientists available to serve each year likely exceeds the number of open editorial positions. While the number of ‘qualified scientists’ is more difficult to quantify, this is also unlikely to be a contributing factor. In 2014 alone, for example, there were over 4200 scientists based in the Global South that were the lead authors of papers in our focal journals – a pool of scientists three times the size of the entire editorial community (Text S1 Table B). Furthermore, 13% of these authors, but only 8% of the editors, were scientists based in middle- and low-income countries, with similar trends for the proportional representation of authors and editors from Africa, the Middle East, Latin America, and the Caribbean (Text S1 Table B). Having said that, we emphasize that it is essential to move beyond proportional representation when thinking of diversity on editorial boards. Why? Because the benefits of diversity continue to accrue as representation increases.

### Why does geographical diversity matter?

Although the increasing Geographic Richness of editors is a positive development, it is dispiriting that Geographic Diversity remains unchanged. Unfortunately, it will remain low until a greater proportion of editors are based outside of the USA and UK. But does a lack of geographic representation – be it at the national, regional, or macroeconomic level – have consequences for the process of evaluating manuscripts that could ultimately limit the scope and direction of research in environmental biology? Put bluntly, do editors and reviewers from high-income regions like the USA or UK have biases – implicit or otherwise – that affect how they evaluate submissions from scientists based in the Global South? Although one journal in our survey found no evidence that reviewer or author nationality influences the likelihood manuscripts are accepted [21,22], this contrasts sharply with the results of prior studies in other STEM fields [23]. There is also compelling evidence that the region in which authors are based affects where their papers are ultimately published and how much they are cited [10,24,25]. In light of these results, and the ample data on how gender and ethnic background influence other aspects of academic evaluation [26], we recommend Editors-in-Chief work to increase the geographic representation on their boards, make editorial board members and referees aware of how biases based on author nationality can affect their editorial judgement, and conduct internal analyses of the potential factors influencing manuscript fate.

Internationalizing editorial boards can also have positive impacts for journals in addition to mitigating possible implicit biases. First, scientists who presume their work will not be judged fairly because of their nationality or where they are based [i.e., the “biased author effect”, 27] may be more likely to submit their manuscripts to journals that have editors representing their region. This both increases the number and scope of submissions a journal receives, and the size and expertise of its reviewer pool. Second, a globally diverse editorial board can serve as an important signal of journal quality and connote prestige [27], especially to those tasked with evaluating individual, institutional, or national scientific productivity [15]. Third, it can enhance the profile and impact of the journal and articles published (to say nothing of justification for editors to demand more support or resources from their publishers). Finally, capacity building is central to the mission of academic societies. By providing editorial opportunities to scholars from emerging scientific regions, society journals can play a pivotal role in achieving this goal.

### Geographic Diversity: Identifying disparities and setting goals

Decades of research have highlighted the positive influence of diversity on scientific research teams [28]. Although we recognize editorial boards do not operate in precisely the same way as workplace teams, we believe that increases in their geographic diversity can similarly enhance the creativity and impact of scholarship published in scientific journals. We reiterate prior calls [16] for journal leadership to, at the very least, strive for editorial boards whose regional distribution of editors mirrors that of authors (Fig 3 and S1 Text Table B, Fig D). However, we also encourage complementing these efforts by including editors based on criteria such as where a journal’s authors work [11,20] and where their expertise is needed [29,30]. Because the size of editorial boards is typically smaller than the number of countries meeting these criteria, we suggest editors attempt to recruit from less-represented countries within a focal region as opportunities arise.

**Fig 3.**
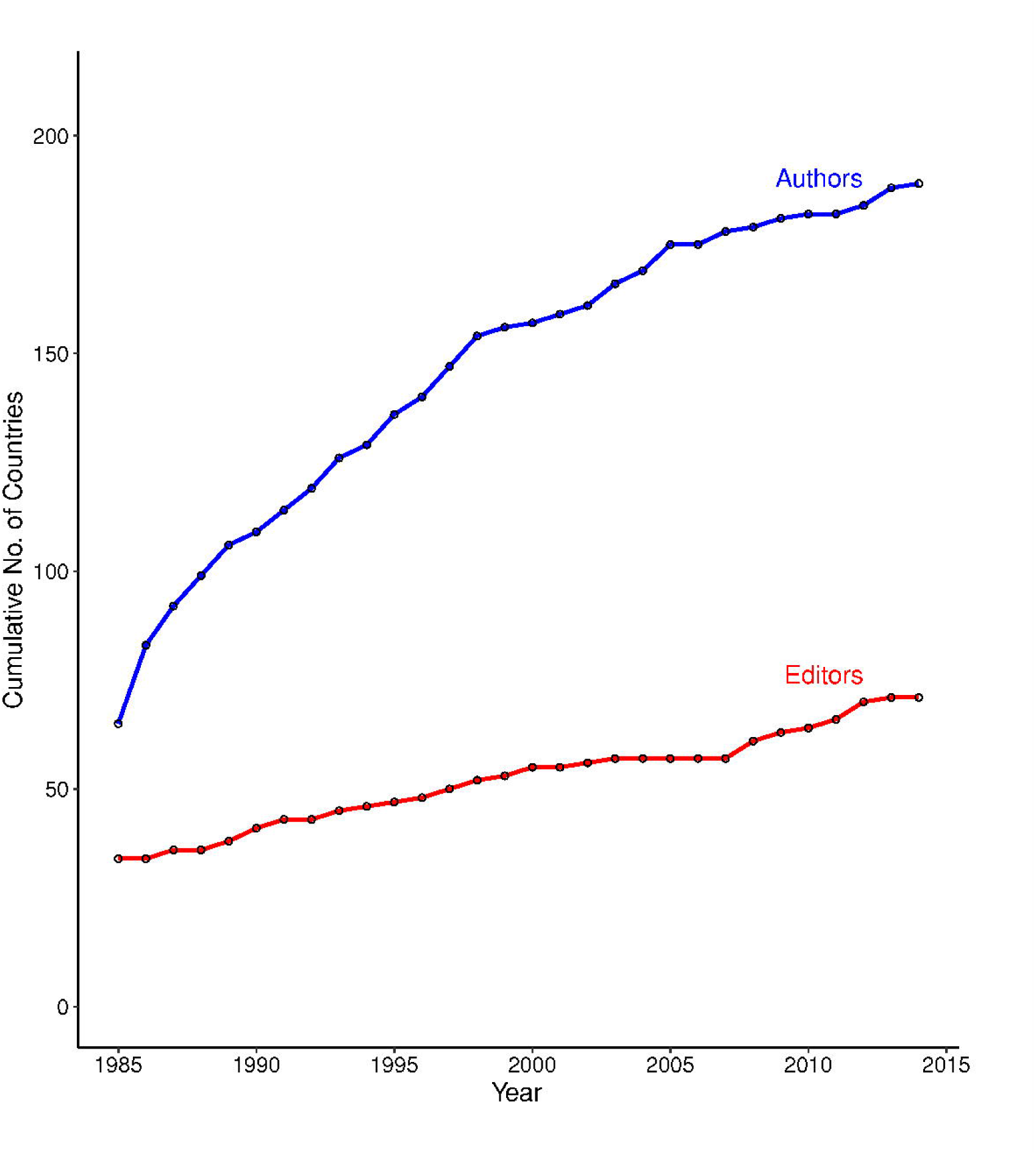
Cumulative Geographic Richness of editors and authors in environmental biology (1985-2014). Rarefaction curves were generated using data on the editorial board membership of 24 environmental biology journals (Table S1) and the institutional addresses of authors publishing in those journals (N = 100,031 articles; S1 Text).

Regardless of the criteria used to identify areas from which to increase representation, however, we efforts must be guided by specific plans and timetables to provide both guidance to editors and hold them accountable for their commitments [31]. Whether such plans underlie the geographic diversity we observed on a few of the editorial boards we reviewed is unknown (Appendix A). Nevertheless, these examples of journals with geographically widespread editors further undermine the frequent argument that it is challenging to find and recruiting board members from the Global South with the requisite academic background, editorial experience, and time to serve. We believe that recruiting these editors is the ethical duty of a journal’s leadership, especially given the impact their presence on the board can have on the global scientific community and the diffusion of the knowledge they create in the service of society. Where to find them? We humbly suggest their large and geographically diverse pool of authors (Fig 3, S1 Text Fig D) is an ideal place to start.

## ACKNOWLEDGEMENTS

We thank Juan Pablo Gomez for helpful discussions and assistance with data collection and T. Bloom, L. Bero, and an anonymous reviewer for comments on the manuscript. This manuscript was completed while EB was Faculty-in-Residence at the University of Florida Marston Science Library.

## REFERENCES

1. Ware M, Mabe M. The STM report: An overview of scientific and scholarly journal publishing. 2015. International Association of Scientific Technical and Medical Publishers.The Hague, The Netherlands. Available at: http://www.stmassoc.org/2015_02_20_STM_Report_2015.pdf

2. Crane D. The gatekeepers of science: Some factors affecting the selection of articles for scientific journals. The American Sociologist. 1967;2(4):195–201.

3. Metz I, Harzing A-W. Gender diversity in editorial boards of management journals. Academy of Management Learning & Education. 2009;8(4):540–57. doi: 10.5465/AMLE.2009.47785472.

4. Garcia-Carpintero E, Granadino B, Plaza LM. The representation of nationalities on the editorial boards of international journals and the promotion of the scientific output of the same countries. Scientometrics. 2010;84(3):799–811. doi: 10.1007/s11192-010-0199-3.

5. Cho AH, Johnson SA, Schuman CE, Adler JM, Gonzalez O, Graves SJ, et al. Women are underrepresented on the editorial boards of journals in environmental biology and natural resource management. PeerJ. 2014;2:e542. doi: 10.7717/peerj.542.

6. Cox TH, Blake S. Managing cultural diversity: Implications for organizational competitiveness. The Executive. 1991:45–56.

7. Mazov NA, Gureev VN. The editorial boards of scientific journals as a subject of scientometric research: a literature review. Scientific and Technical Information Processing. 2016;43(3):144–53. doi: 10.3103/s0147688216030035.

8. Fox CW, Burns CS, Meyer JA, Thompson K. Editor and reviewer gender influence the peer review process but not peer review outcomes at an ecology journal. Functional Ecology. 2016;30(1):140–53. doi: 10.1111/1365-2435.12529.

9. Holmgren M, Schnitzer SA. Science on the rise in developing countries. PLoS Biol. 2004;2(1):e1. doi: 10.1371/journal.pbio.0020001.

10. Smith MJ, Weinberger C, Bruna EM, Allesina S. The scientific impact of nations: Journal placement and citation performance. PLoS ONE. 2014;9(10):e109195. doi: 10.1371/journal.pone.0109195.

11. Mammides C, Goodale UM, Corlett RT, Chen J, Bawa KS, Hariya H, et al. Increasing geographic diversity in the international conservation literature: A stalled process? Biological Conservation. 2016;198:78–83. doi: 10.1016/j.biocon.2016.03.030.

12. Meneghini R, Packer AL, Nassi-Calo L. Articles by Latin American authors in prestigious journals have fewer citations. PLoS ONE. 2008;3(11). doi: 10.1371/journal.pone.0003804.

13. Bakker P, Rigter H. Editors of medical journals - who and from where. Scientometrics. 1985;7(1-2):11–22. doi: 10.1007/bf02020137.

14. Braun T, Bujdoso E. Gatekeeping patterns in the publication of analytical-chemistry research. Talanta. 1983;30(3):161–7. doi: 10.1016/0039-9140(83)80043-5.

15. Nisonger TE. The relationship between international editorial board composition and citation measures in political science, business, and genetics journals. Scientometrics. 2002;54(2):257–68. doi: 10.1023/A:1016065929026.

16. Livingston G, Waring B, Pacheco LF, Buchori D, Jiang YX, Gilbert L, et al. Perspectives on the global disparity in ecological science. Bioscience. 2016;66(2):147–55. doi: 10.1093/biosci/biv175.

17. Independent Commission on International Development Issues. North-South: a programme for survival: report of the Independent Commission on International Development Issues. Cabridge, MA: MIT Press; 1980.

18. Habel JC, Lens L, Eggermont H, Githiru M, Mulwa RK, Shauri HS, et al. More topics from the tropics: additional thoughts to Mammides et al. Biodiversity and Conservation. 2017;26(1):237–41. doi: 10.1007/s10531-016-1236-1.

19. UNESCO. Number of Researchers per million inhabitants by country, United Nations Educational, Scientific and Cultural Organisation (UNESCO) Institute for Statistics. 2011. Available at:

20. Stocks G, Seales L, Paniagua F, Maehr E, Bruna EM. The geographical and institutional distribution of ecological research in the tropics. Biotropica. 2008;40(4):397–404. doi: 10.1111/j.1744-7429.2007.00393.x.

21. Campos-Arceiz A, Primack RB, Koh LP. Reviewer recommendations and editors' decisions for a conservation journal: Is it just a crapshoot? And do Chinese authors get a fair shot? Biological Conservation. 2015;186:22–7. doi: 10.1016/j.biocon.2015.02.025.

22. Primack RB, Ellwood E, Miller-Rushing AJ, Marrs R, Mulligan A. Do gender, nationality, or academic age affect review decisions? An analysis of submissions to the journal Biological Conservation. Biological Conservation. 2009;142(11):2415–8. doi: 10.1016/j.biocon.2009.06.021.

23. Lee CJ, Sugimoto CR, Zhang G, Cronin B. Bias in peer review. Journal of the American Society for Information Science and Technology. 2013;64(1):2–17. doi: 10.1002/asi.22784.

24. Mori AS, Qian SH, Tatsumi S. Academic inequality through the lens of community ecology: a meta-analysis. PeerJ. 2015;3(3):e1457. doi: 10.7717/peerj.1457.

25. Meijaard E, Cardillo M, Meijaard EM, Possingham HP. Geographic bias in citation rates of conservation research. Conserv Biol. 2015;29(3):920–5. doi: 10.1111/cobi.12489.

26. Menges RJ, Exum WH. Barriers to the progress of women and minority faculty. The Journal of Higher Education. 1983;54(2):123–44. doi: 10.1080/00221546.1983.11778167.

27. Pearson C, Mullen R, Thomason W, Phillips S. Associate editor's role in helping authors and upholding journal standards. Agronomy Journal. 2006;98(3):417–22. doi: 10.2134/agronj2005.0296.

28. Liao C. How to improve research quality? Examining the impacts of collaboration intensity and member diversity in collaboration networks. Scientometrics. 2011;86(3):747–61. doi: 10.1007/s11192-010-0309-2.

29. Campos-Arceiz A, Primack RB, Miller-Rushing AJ, Maron M. Striking underrepresentation of biodiversity-rich regions among editors of conservation journals. Biol Conserv. 2017. doi: doi.org/10.1016/j.biocon.2017.07.028.

30. Karlsson S. The North-South knowledge divide: consequences for global environmental governance. Strengthening Global Environmental Governance: Options and Opportunities: Yale School of Forestry & Environmental Studies; 2002. p. 1–24.

31. Calver M, Wardell-Johnson G, Bradley S, Taplin R. What makes a journal international? A case study using conservation biology journals. Scientometrics. 2010;85(2):387–400. doi: 10.1007/s11192-010-0273-x.

